# Shared Transcriptomic Signatures of Inflammaging Among Diverse Strains of *Drosophila melanogaster*

**DOI:** 10.1101/2024.01.17.576061

**Authors:** Sabrina Isabel Perna, Weihao Tang, Sydney Danielle Blimbaum, Andrew Li, Lei Zhou

## Abstract

**Background:** A prominent hallmark of aging is inflammaging—the increased expression of innate immune genes without identifiable infection. Model organisms with shorter lifespans, such as the fruit fly, provide an essential platform for probing the mechanisms of inflammaging. Multiple groups have reported that, like mammalian models, old flies have significantly higher levels of expression of anti-microbial peptide genes. However, whether some of these genes—or any others—can serve as reliable markers for assessing and comparing inflammaging in different strains remains unclear.

**Methods and Results:** We compared RNA-Seq datasets generated by different groups. Although the fly strains used in these studies differ significantly, we found that they share a core group of genes with strong aging-associated expression. In addition to anti-microbial peptide genes, we identified other genes that have prominently increased expression in old flies, especially *SPH93*. We further showed that machine learning models can be used to predict the “inflammatory age” of the fruit fly.

**Conclusion:** A core group of genes may serve as markers for studying inflammaging in *Drosophila*. RNA-Seq profiles, in combination with machine-learning models, can be applied to measure the acceleration or deceleration of inflammaging.

## Introduction

Innate immunity and inflammation are essential for defense against infection. However, inflammation may linger in older animals, often without identifiable infection—a phenomenon referred to as *inflammaging*. Inflammaging is implicated in many age-related diseases, such as arthritis, neurodegenerative diseases, and cancers [1]. One sign of inflammaging is provided by transcriptomic studies comparing the gene-expression profiles of old vs. young animals [2,3]. Inflammatory genes are significantly overexpressed as a result of aging in all mammalian species that have been subjected to transcriptomic monitoring [4,5].

Many competing theories and models have been proposed to explain the mechanisms underlying inflammaging (reviewed by Ferrucci and Fabbri [6]). However, progress has been hindered by the inherent difficulty of studying inflammaging in mammalian models with long lifespans. Fruit flies may be a promising alternative model since they share important innate immune pathways with mammals but have a lifespan of just 50–70 days [7–9]. Most importantly, there is evidence that the fundamental mechanisms of inflammaging are well-conserved between fruit flies and mammals. For instance, inflammaging in fruit flies is significantly suppressed by the human serum protein alpha antitrypsin (hAAT), which modulates the same innate immune pathway in fruit flies and human senescent cells [10].

Quantitative measurement is essential for mechanistic studies of inflammaging. In humans, biological age can be assessed using DNA methylation levels [11], and inflammatory aging can be monitored by cytokine levels or gene expression in the blood [12,13]. In the fruit fly, in contrast, gene expression seems to be the only practical option. In parallel to what has been observed in mammalian models, transcriptomic studies in older flies have revealed significant upregulation of innate immune genes, mostly anti-microbial peptide (AMP) genes [14,15]. RNA-Seq analysis in Drosophila also revealed that immune-defense genes are overrepresented in genes upregulated with age [16]. Yet it remains unclear which AMP genes, or others, can serve as reliable markers for assessing inflammation in the fruit fly.

In this study, we utilized RNA-sequencing datasets to identify potential hallmarks of inflammaging that are shared across various strains of the fruit fly. Our analysis showed that different marker genes should be considered to assess inflammaging in males vs. females. In addition, we found that whole-body RNA samples are more reliable than dissected samples in assessing inflammaging. We propose that RNA-Seq profiles, combined with machine-learning models, can be used to measure the *Drosophila* inflammatory age (DiAge) for mechanistic studies.

## Methods Summary

This study used a few novel methods, summarized in brief here to help the reader interpret the results. A complete description can be found in the Detailed Methods section.

### AGE-Index

To highlight genes whose expression levels are significantly increased in older fruit flies, we assigned the <u>a</u>ging-associated <u>g</u>ene <u>e</u>xpression index (AGE-Index) for each gene as the ratio of its expression level in the old stage (30+ days after eclosion) divided by that in the young stage (3–10 days after eclosion). Since the expression levels of certain innate immune genes differed significantly between males and females, we opted to calculate the AGE-Index separately for each sex. The AGE-Index values are conceptually equivalent to the fold change that is often measured by QPCR analysis for individual genes.

Four datasets were used for calculating AGE-Index for whole-body samples (Table 1), including two well-known datasets from the public domain and two RNA-Seq datasets generated by us. To avoid any bias introduced by our mapping and quantification process, for the public datasets (*modENCODE*, *TM*), we used the expression values calculated by the original data producers to derive the AGE-Index values. For RNA-Seq data generated by us (*Oregon-R*, *w1118*), transcripts per million (TPM) values were obtained using Salmon [17] before calculating the AGE-Index.

**Table 1.** A list of whole-body RNA-Seq datasets used for identifying inflammaging patterns.

| *Codename* | *Strain(s)* | *Ages assayed* | *Culture condition* | *RNA extraction* | *Sequencing platform* | *Publication* | *Expression Profile data* |
| --- | --- | --- | --- | --- | --- | --- | --- |
| modENCODE | Isogenic *y<sup>1</sup>*; *cn bw<sup>1</sup>sp<sup>1</sup>* | D5, D30 | 25 °C?<br>Standard food? | Trizol, Qiagen column purification. | 454, Illumina, | Graveley et al. 2011 | PRJNA75285; “gene_rpkm_report_fb_2021_05.tsv” |
| TM | 5 base (B) lines, 5 (O) lines selected for postponed senescence. | Week1, Week5 | 25 °C;<br>Cornmeal-molasses-gar medium. | Trizol (Life Technologies) and RNAeasy mini (Qiagen) | Illumina (TrueSeq / Hiseq2500) | Carnes et al. 2015. | PRJNA286855; S4_Table of the publication |
| Oregon-R | *Oregon-R* (RRID: BDSC_6363) | D5, D30 | 25 °C;<br>Bloomington recipe food. | Trizol | Illumina | This study | TBA |
| W1118 | *w1118* (RRID: BDSC_5905) | D5, D30, D45 | 25 °C;<br>Bloomington recipe food. | Trizol | Illumina | This study | TBA |

### Immune-defense gene list

To focus our attention on the genes involved in immunity and defense against pathogens, we compiled a list of genes associated with GO:0002376, “immune system process,” or GO:0098542, “defense response to other organisms”. The Immune-defense (ImmDef) gene list has 504 genes marked as solid red dots in AGE-Index plots.

### The machine learning model for predicting DiAge

To test if RNA-Seq profiling can be used to determine the biological age of the fruit fly, we built elastic-net regression models using RNA-Seq profile of wild type flies of various age. Most of these were samples used as controls for various studies. The dataset was compiled by downloading the raw reads and obtaining TPM using Salmon. Inspired by the work of Horvath [11], we transformed the chronological age of the samples to T_age = (age − 1)/5. For each gene, its expression level in a given sample x was transformed to (TPMx − TPMmin) / (TPMmax − TPMmin). The matrix containing T_age and min-max normalized gene expression values was then used to train the elastic-net model using the R package glmnet [18].

## Results

### Shared inflammaging patterns among diverse strains of *D. melanogaster* at the whole-body level

To gain a comprehensive view of genes with increased expression in old animals, we plotted the female and male AGE-Index values for each gene along the x-axis and y-axis, respectively (Figure 1). The overall pictures vary considerably among the different strains, which were assessed by different labs spanning more than a decade (Table 1). Yet it is clear that at the whole-body level in all the strains, genes involved in the immune response (GO:0002376) or defense response (GO:0098542) are prominent among the genes with high AGE-Index values (Figure 1, red dots).

**Figure 1.**
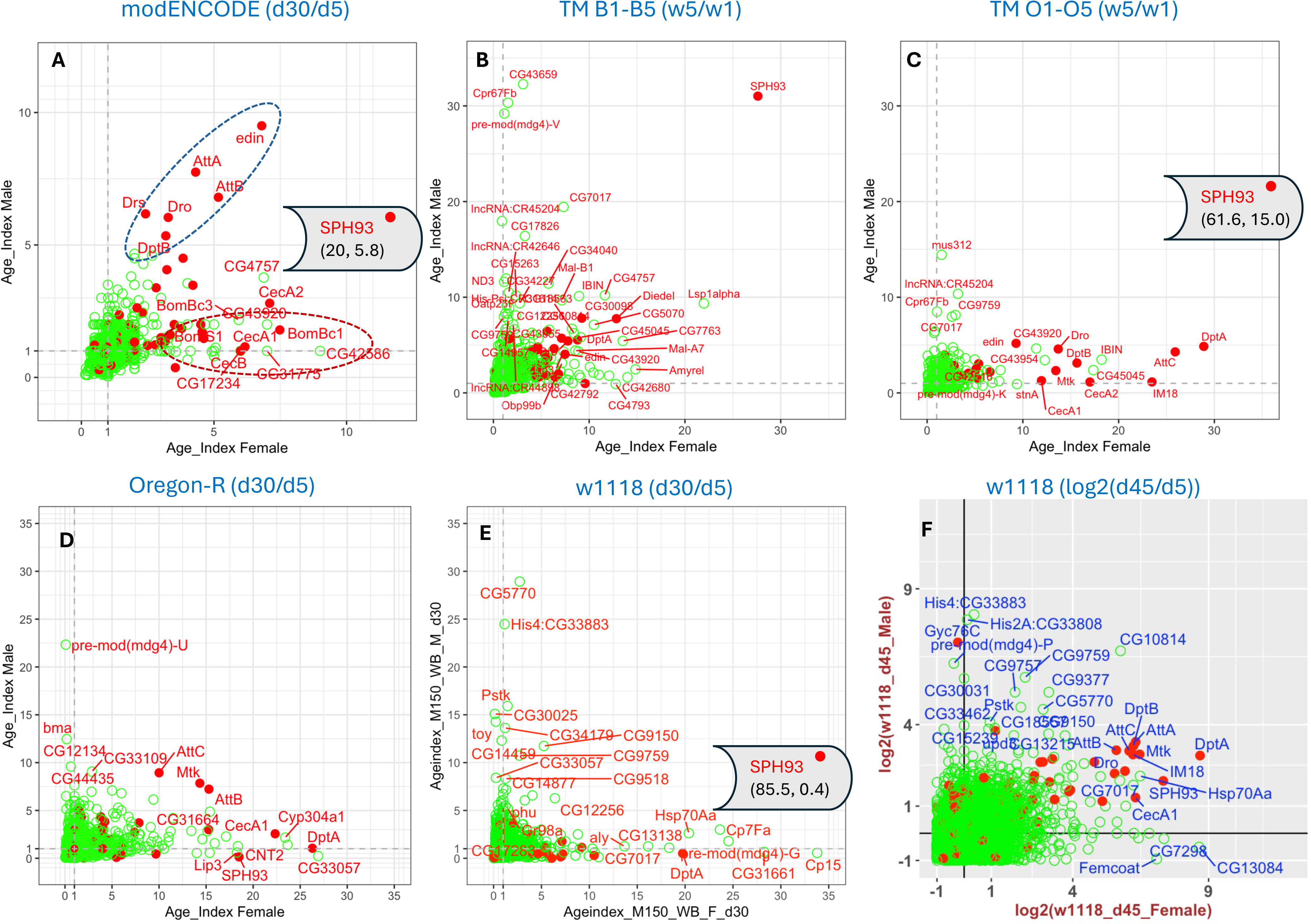
AGE-Index based on whole-body RNA-Seq datasets. Dot plot of genes based on the AGE-Index for females (x-axis) and males (y-axis). Names for genes with AGE-Index greater than 5 in either sex were included in the plot. Genes involved in immune (GO:0002376) or defense response (GO:0098542) processes were marked as solid red dots. All others were represented as green circles. Genes with decreased expression in old animals were not included in the plot. (A) AGE-Index based on the ModENCODE developmental stage dataset. One gene, SPH93, falls out of the plot range. Its respective AGE-Index values in females and males are indicated in the parenthesis, respectively. (B) AGE-Index plot for the average of five B (base) lines included in the dataset generated by Garnes et al. (C) AGE-Index plot for the average of five O lines selected for postponed senescence. One gene falling out of the plot range is SPH93, its AGE-Index values for females and males are indicated as (x, y) coordinates in the parenthesis. (D) AGE-Index plot for Oregon-R at day 30. (E,F) AGE-Index plot for w1118 at day 30 (E) and day 45 (F). The plot on F is log2 transformed AGE-Index values. All others are plotted as untransformed. .

For the *modENCODE* developmental-stage dataset [19], sixteen genes have an AGE-Index value greater than 5 in either sex. Five of those only have a CG (computer-generated) number. All the named genes are annotated as participating in immune- or defense-response functions (Figure 1A). Three genes (*SPH93*, *edin*, *AttB*) have an AGE-Index value greater than 5 in both sexes. *SPH93* had such a dramatic change between days 5 and 30 that it had to be excluded from the plotting to allow discernment of the other genes. The whole-range plot that includes *SPH93* can be found in SF2.

For the second *TM* dataset, the average AGE-Index values were calculated separately for the five B (base) lines (Figure 1B) and the five O lines (Figure 1C) since the expression of innate immune genes was found to differ between the two groups [16].

Two features stand out in this dataset. First, the magnitude of the increase of innate immune genes is more prominent for the O lines than the B lines. Second, like what we saw with the *modENCODE* dataset, *SPH93* has by far the highest age-associated increase of expression in week-5 vs. week-1 flies (Figure 1B & C).

For the *Oregon-R* dataset, we processed two replicates of day-5 and day-30 RNA samples from both sexes and calculated the AGE-Index value based on the TPM (Figure 1D). In addition, genes with significant changes in expression were identified using CuffDiff and DESeq2 (S4) [20]. The immune-defense genes that show a prominent increase in old age in this strain overlap significantly with those identified in the public datasets (Table 2).

**Table 2.** Top 12 inflammaging marker genes based on their *AGE_Index* value. Genes shared by all datasets are in bold.

| Rank | Male |  |  |  |  | Female |  |  |  |  |
| --- | --- | --- | --- | --- | --- | --- | --- | --- | --- | --- |
|  | *ModENCODE* | *TM-B lines* | *TM-O lines* | *Oregon-R* | *w1118* | *ModENCODE* | *TM-B lines* | *TM-O lines* | *Oregon-R* | *w1118* |
| 1 | edin | SPH93 | SPH93 | **AttC** | Gyc76C | **SPH93** | **SPH93** | **SPH93** | AttA | **DptA** |
| 2 | **AttA** | CG30098 | edin | Mtk | upd3 | BomBc1 | Diedel | **DptA** | **DptA** | **SPH93** |
| 3 | **AttB** | Diedel | DptA | **AttB** | **AttA** | CecA2 | lncRNA:CR43855 | AttC | CecA1 | Mtk |
| 4 | Drs | **AttA** | Dro | **AttA** | DptB | edin | CG30098 | IM18 | **SPH93** | AttA |
| 5 | Dro | CG43055 | **AttC** | Drsl2 | **AttB** | CecA1 | **DptA** | CecA2 | AttB | CecA1 |
| 6 | SPH93 | upd3 | Def | CG43055 | **AttC** | CecB | Dro | DptB | DptB | DptB |
| 7 | DptB | DptA | DptB | LysX | Mtk | AttB | edin | Dro | Mtk | IM18 |
| 8 | **AttC** | Dro | **AttA** | PGRP-SB1 | IM18 | GNBP-like3 | CG43055 | Mtk | AttC | AttC |
| 9 | Def | **AttB** | upd3 | BaraA1 | DptA | Dso1 | IM18 | CecA1 | CecC | Dro |
| 10 | DptA | Def | **AttB** | IM18 | CecB | AttD | Toll-4 | edin | IM18 | AttB |
| 11 | Mtk | **AttC** | Listericin | CG12780 | BomBc1 | AttA | AttC | CecC | Dro | CecA2 |
| 12 | CecA2 | AttD | CG43055 | Diedel | AttD | **DptA** | Mtk | AttA | Vago | Drsl2 |

Finally, for the *w1118* strain, on day 30, only a few genes, including *SPH93* and *DptA*, show significant increases in expression in the females but not the males (Figure 1E, S5). Because of this, we also analyzed *w1118* samples on day 45 and found that the expression of many innate immune genes was significantly higher in both sexes (Figure 1F). For visualization purposes, the day-45 AGE-Index values are log_2-_transformed; the non-transformed plot can be found in SF2.

Very few immune-defense genes show decreased expression in old age. To reveal genes whose expression is negatively associated with aging, we plotted log_2_(AGE-Index) for all immune-defense genes (SF2). In the three datasets we produced, PPO1 and PPO2 were significantly decreased in old ages (SF2). However, this was not seen for the modENCODE or the TM datasets. In contrast, the shared pattern of immune-defense genes positively correlating with aging is apparent (Figure1). A ranking of the top 12 genes with the highest AGE-Index values is shown in Table 2. Across all datasets, the three Attacin genes (*AttA*, *AttB*, *AttC*) were reliably present for males, while *SPH93* and *DptA* were reliably present for females.

### Female-skewed inflammaging pattern

The plotting of female vs. male AGE-Index values revealed sexual differences in inflammaging at the whole-body level (Figure 1). In the *modENCODE* dataset, for example, immune-defense genes with high AGE-index values can be roughly divided into two groups. The first has high AGE-Index values in both sexes, which we call the shared inflammaging pattern (Figure 1A, dashed blue circle). The second group consists of genes whose AGE-Index values in females is high but in males is either close to 1 or much lower than the female values. We refer to this as the female-skewed inflammaging pattern (dashed red circle, Figure 1A).

For many of the inflammaging markers we identified, the *TM-O* average (Figure 1C) shows a prominent female-skewed inflammaging pattern, apparent for each of the five *TM-O* lines (SF3). In contrast, the *TM-B* average had few inflammaging markers that fell into this pattern (Figure 1B), and only one of the five *TM-B* lines had any genes in this pattern (B2, SF3). Since the same research lab processed the ten TM lines in parallel, the differences in patterns among these lines are most likely due to genetic and epigenetic variations rather than differences in the growing environment and sample processing. Thus, we focused on this dataset to understand what underlies the female-skewed inflammaging pattern.

We examined the expression levels of five inflammaging markers at the young and old stages in both females and males (Figure 2). This revealed two reasons why AGE-Index values were higher in females than males in the *TM-O* lines. The first is suppressed expression of the immune-defense gene in young *TM-O* females. For instance, the inflammaging pattern of *DptA* in the *TM-B* lines is not female-skewed, with average female and male AGE-Index values of 8.6 and 5, respectively (Figure 1B, SF2). In the *TM-O* lines, its inflammaging pattern is highly skewed with female and male AGE-Index values of 24.8 and 4.7, respectively (Figure 1C, SF2). The very high female AGE-Index value of *DptA* in *TM-O* lines is mostly due to the significantly decreased expression of *DptA* in *TM-O* females at week 1 (p < 0.01, Figure 2A). At week 5, *TM-O* females have slightly lower expression levels of *DptA* than *TM-B* females, yet the AGE-Index values of *DptA* are much higher due to the extremely low reference point of the week-1 level. The same is true for *AttC* (Figure 2B) and several other genes that showed only female-skewed patterns in *TM-O* but not *TM-B* lines.

**Figure 2.**
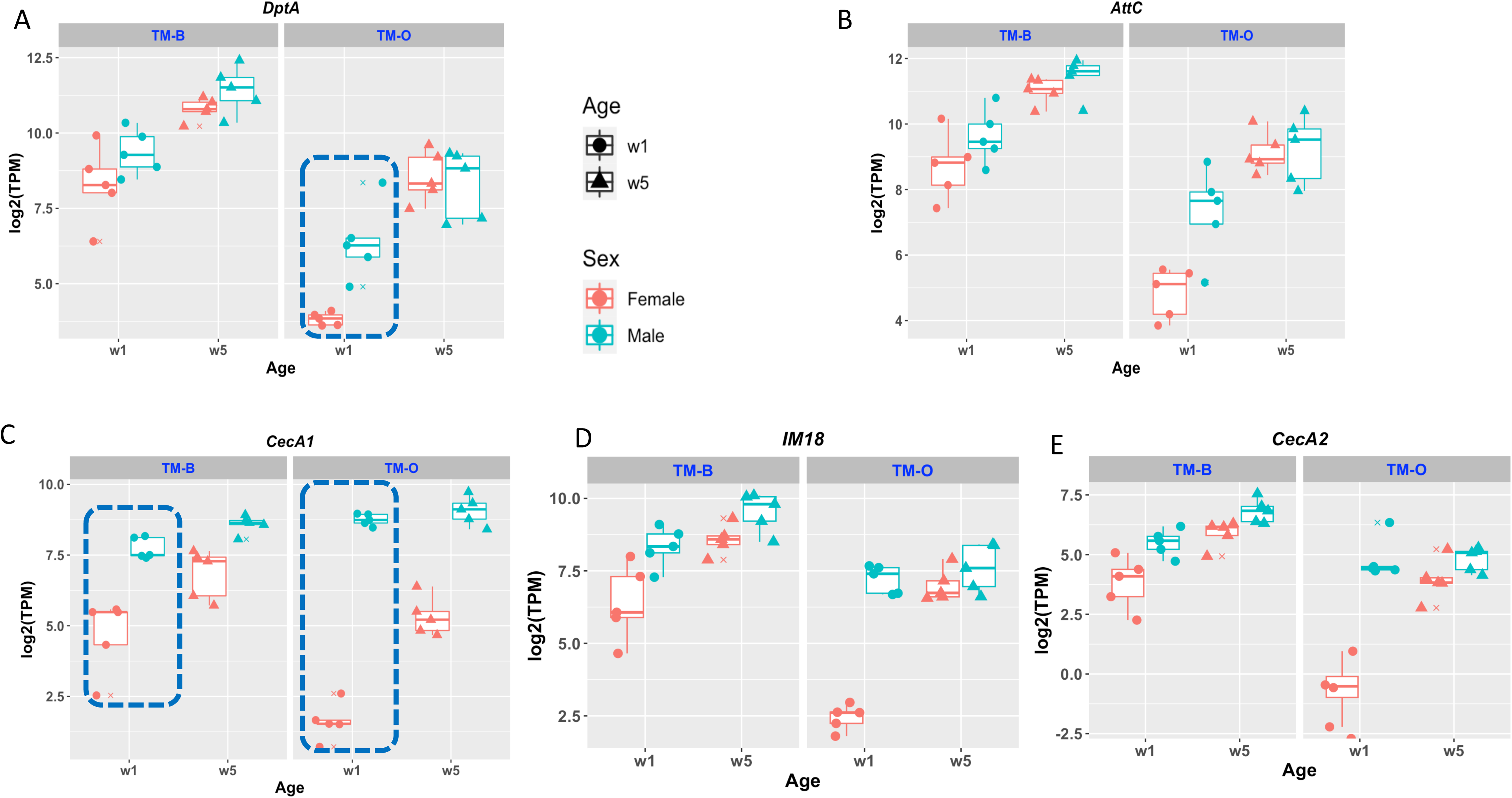
Expression of the inflammaging marker genes in the TM-B and TM-O lines. Each panel depicts the expression levels of one inflammaging marker gene in the TM dataset. The five TM-B lines and Five TM-O lines are grouped into different facets. Each data point reflects the average of two replicate RNA-Seq experiments.

The second reason for the female-skewed inflammaging pattern is revealed by comparing the expression levels of *CecA1* (Figure 2C), a gene with a consistent female-skewed pattern across both *TM-B* and *TM-O* lines. Its average AGE-Index values for females and males were 5.30 & 1.83 in the *TM-B* line and 10.25 & 1.28 in the *TM-O* line, respectively (Figures 1B & 1C, SF2). However, this sexual difference in AGE-Index appears mostly due to the much higher levels of *CecA1* expression in young males in all lines (dotted circles). It is known that *CecA1* is expressed at high levels in the male reproductive tract [21], which we verified using transgenic fly strains carrying *CecA1*-GFP reporter. The very high expression of *CecA1* in the male reproductive organs significantly increases the denominator for calculating whole-body AGE-Index values, thus masking potential inflammaging patterns of this gene in other organs, such as the head (see the following section). The other gene that consistently demonstrated the female-skewed inflammaging pattern is *IM18 (Figure 2D)*, which is highly expressed in male germline cells and testis [22]. However, suppression of *IM18* is also apparent in the *TM-O* females compared to *TM-B* females. This has been observed for other genes such as *CecA2* (Figure 2E).

### Expression of innate immune genes in head and fat body

To avoid the compounding effect of sexual-organ-specific expression, we next compared the expression values of innate immune genes in the heads and fat bodies of male and female flies (Figure 3). We found that the sexually dimorphic expression pattern of some immune-defense genes is organ-specific and age-specific.

**Figure 3.**
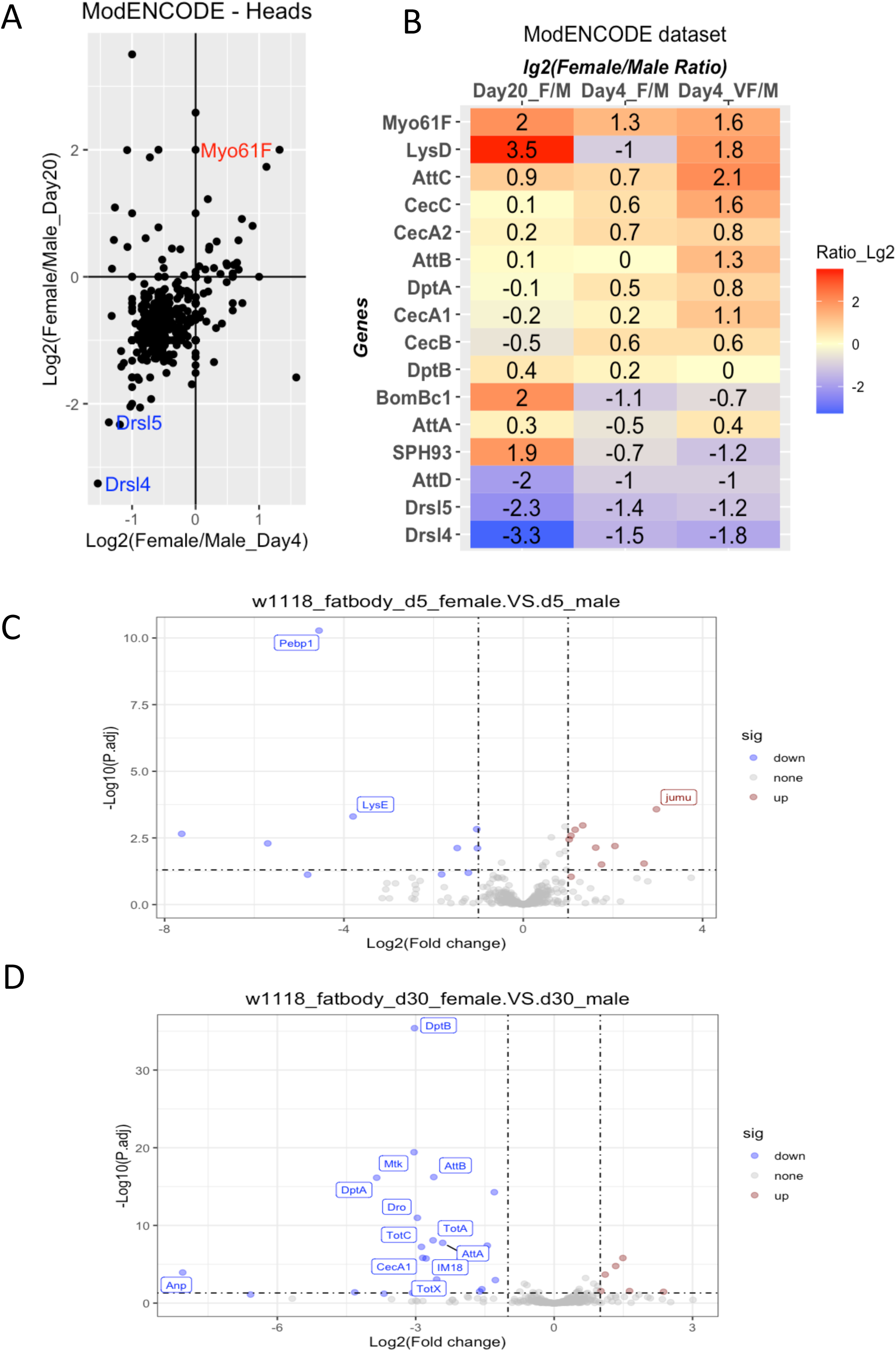
Immune and defense genes display sexual dimorphic expression patterns. A & B. Expression of immune & defense genes in the heads of male and female flies based on the ModENCODE dataset. F-female, VF-virgin female, M-male. C& D. Volcano plot of gene expression for w1118 female vs. male fat bodies at day 5 (C) and day 30 (D). P values obtained by DESeq2 analysis of RNA-Seq profiles of 3 biological repeats.

The *modENCODE* dataset has expression profiles for male and female heads obtained on day 4 and day 20. Figure 3A shows the log_2_-transformed female-to-male ratio of all genes involved in immune and defense processes. The comparison revealed that for the majority of genes involved in these two processes, their expression levels are slightly lower in female heads than in male heads on both days 4 and 20. One gene (*Myo61F*) showed female-biased expression, and two (*Drsl5* and *Drsl4*) showed male-biased expression. However, many genes have biased expression levels only in one stage but not the other. Notably, most of the major inflammaging markers we identified, such as *DptA*, *SPH93*, *AttA*, and *AttB*, do not show significant sex-biased expression in this dataset (Figure 3B).

Besides the head expression profile in the *modENCODE* dataset, we could not find datasets suitable to assess the potential sexual dimorphism of innate immune gene expression in other tissues or organs. Thus, we determined the expression profile of the fat bodies for both sexes on days 5 and 30. We found that on day 5, the expression profile of immune-defense genes is largely similar between male and female fat bodies. However, by day 30, there is a significant difference between the fat bodies of the males vs. females (Figure 3C & 3D). Many of the inflammaging markers, such as *DptB* and *AttB*, have significantly lower expression in the female fat body than in the male fat body.

### Distinct dynamics of inflammaging in organs and body parts

Our analysis of whole-body RNA-Seq indicated that the dynamics of inflammaging vary among different strains. We next asked whether inflammaging in different organs or body parts follows the same or different dynamics and sought to compare organ-specific inflammaging patterns between *white Dahomey* (*wDah*) and *w1118*.

The female *wDah* dataset (GSE130158) allowed us to compare levels of inflammaging in four different organs or body parts (Figure 4A–D). For each gene involved in the immune-defense processes, its day-30 and day-50 AGE-Index values were plotted along the y-axis and x-axis, respectively (Figure 4A–D). For the brain and thorax, AGE-Index values of inflammatory markers such as *SPH93* are close to 1 on day 30, indicating the expression level is about the same between days 10 and 30. By day 50, however, the levels of *SPH93* in the head and thorax datasets had increased more than seven-fold (Figures 4A & 4B). In contrast, for the fat body, the AGE-Index values of inflammatory marker genes on days 30 and 50 were about the same (Figure 4C). This indicates that, at least for this particular strain, the dynamics of inflammaging are different among the organs or body parts. The fat body showed clear signs of inflammation on day 30. In the brain and thorax, the increased levels of innate immune genes were minimal on day 30 but reached a much higher level on day 50. For the gut, however, only one gene (*upd1*) had an AGE-Index value higher than 5 on day 50 (Figure 4D).

**Figure 4.**
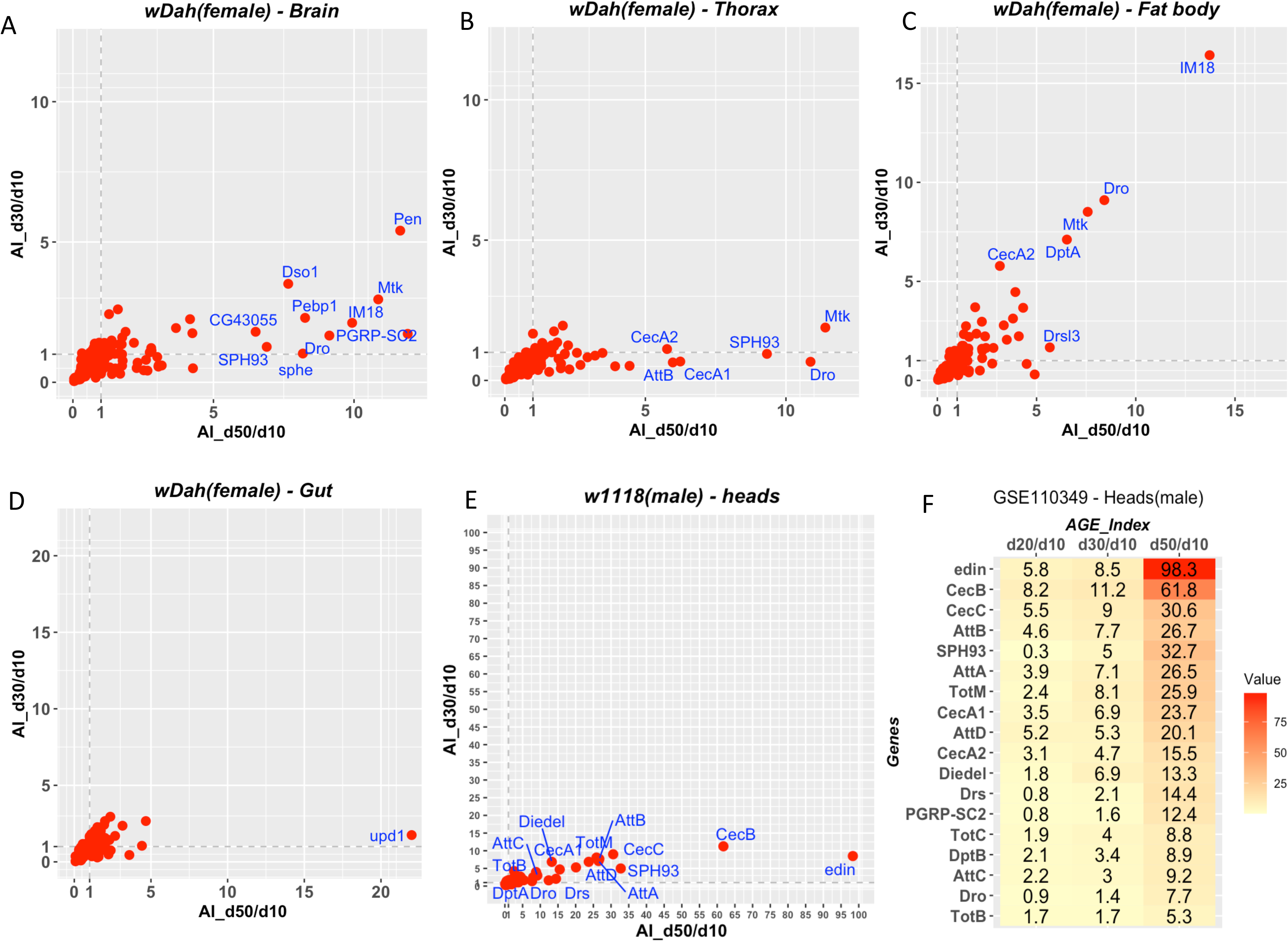
Inflammaging patterns for body parts and organs. A-D, Plotting of AGE-Index values for Brain (A), Thorax (B), fat body (C), and Gut (D) based on the dataset of GSE130158. The expression profiles were generated with females of the white Dahomey (wDah) strain. E-F, dynamics of inflammaging of w1118 male heads based on GSE110349. E, plotting of AGE-Index values for day-30/day-10 (y axis), and day-50/day-10 (x-axis). F, heatmap for day20/day10, day30/day10, day50/day10. Genes names are only plotted for those having an AGE-Index value greater than five.

We next sought to compare organ-specific patterns between this strain and the *w1118* strain. For the head, we used the data provided in GSE110349 [23]. Similar to what was observed for *wDah*, inflammation in the head of *w1118* was much more prominent on day 50 than day 30 (Figure 4E). Several genes, such as *SPH93* and *Dro*, showed similar patterns in the heads of these two strains. Interestingly, this dataset also revealed that *CecA1* and *CecA2* levels in male heads increase significantly as the animal ages (Figure 4F).

For the fat body of *w1118*, we generated our own dataset for both females and males on days 5, 30, and 45 (Figure 5). Unlike what we saw with w*Dah*, there is a significant progression of inflammation between days 30 and 45 (Figure 5 B vs. A). The AGE-Index values were much higher for the *w1118* strain and had to be log2-transformed before plotting.

**Figure 5.**
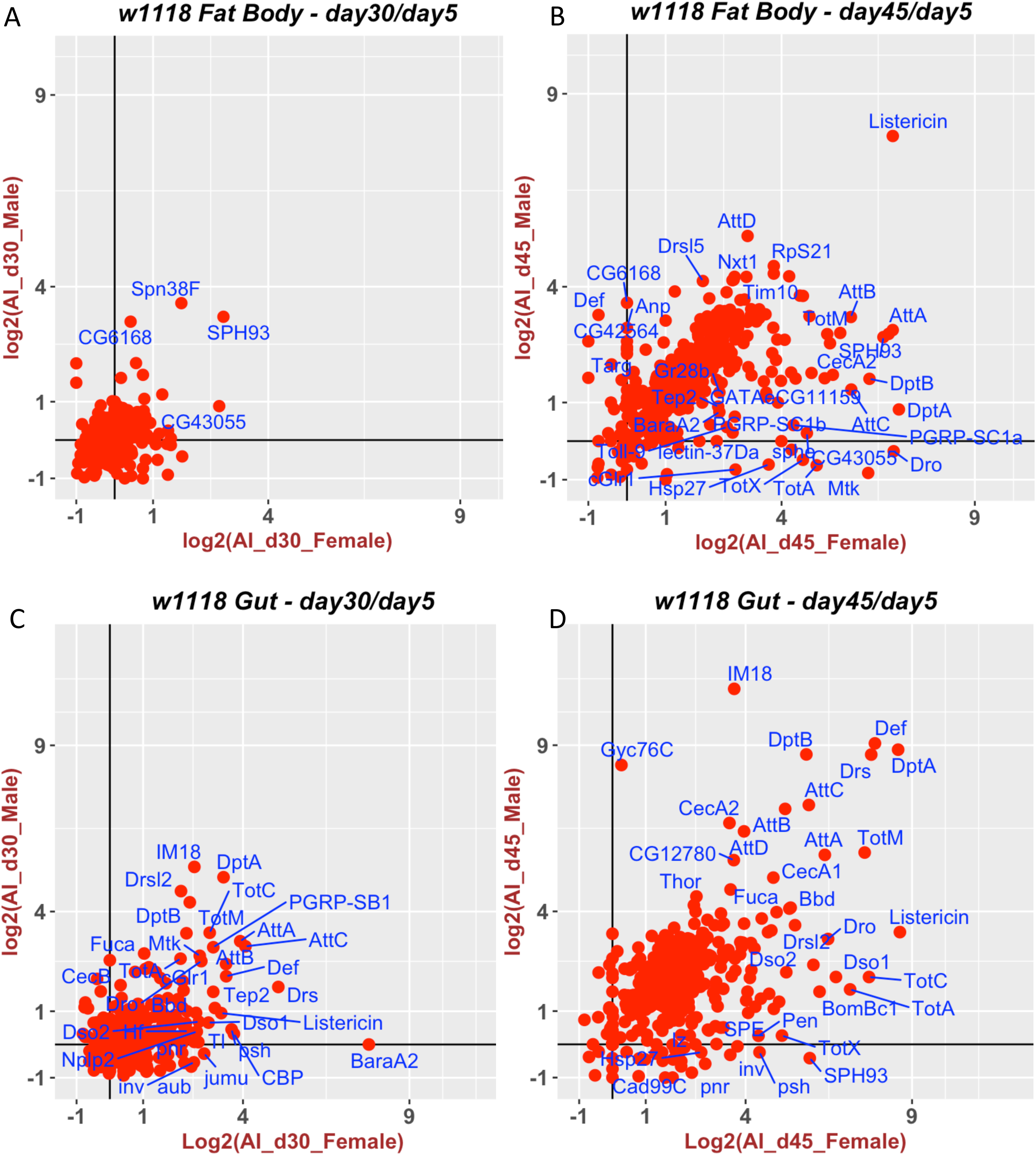
Dynamics of inflammaging for the fat body and gut of the w1118 strain. AGE-Index values for fat bodies (A&B) and whole gut (C&D) in w1118 males and females were calculated based on RNA-Seq samples collected at day 30 (A, C) and day 45 (B, D) and using the day 5 expression level as the reference. The AGE-Index values were plotted as Log2 transformed, with male values on the y-axis and female values along the x-axis. Genes names are only plotted for those having an AGE-Index value greater than five.

The difference in gut inflammaging between these two strains was even more dramatic. Many genes had an AGE-Index value higher than 5 in the gut of *w1118* on day 30 (Figure 5C). This progressed to much higher levels on day 45 (Figure 5D).

In summary, it appears the inflammaging pattern in the head is more consistent between these two strains than that in the fat body or gut.

### Prediction of “inflammatory age” (DiAge) with a machine-learning model

Drawing on various studies (S7), we compiled a machine-learning dataset based on RNA-Seq profiles of the whole body, heads, or body without heads. The age distribution of this learning set is shown in Figure 6A. It is skewed towards young age, with an average age of ∼17 days and median age of just 9.5 days. The sample set was randomly split 0.7:0.3 for training and testing. The best model achieved a mean absolute error (MAE) value of 0.503, corresponding to ∼2.62 days (Figure 6C). We then limited this machine-learning approach to only males. Although the number of male samples is less than half of the total samples, the resulting regression model achieved a mean MAE of 0.321 (∼1.57 days).

**Figure 6.**
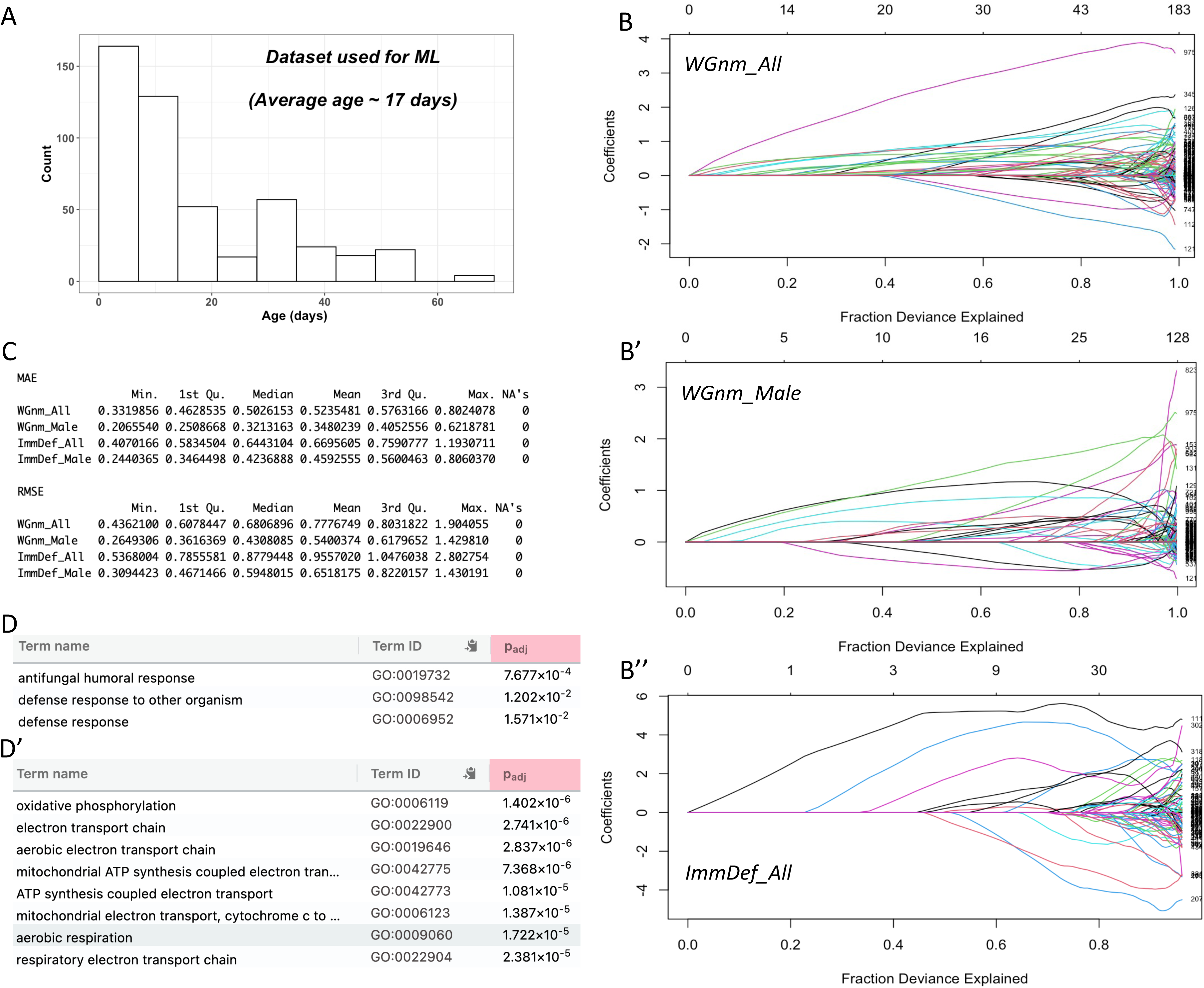
Elastic-net model for inflammatory age prediction. A. Age distribution of samples used to build and test models. B-B”, Plot of coefficients, bottom axis – fraction of deviance explained, top axis – number of variables involved. B-model built with whole genome expression profiles of all samples (WGnm_All). B’ – model built with whole genome profiles of male samples (WGnm_Male). B’’-model built with only genes involved in immune or defense processes (ImmDef_All). C. Assessment of models via resampling cross-validation. MAE-Mean Absolute Error, RMSE-Root Mean Squared Error. D-D’. Gene Ontology (GO) biological process (BP) terms enriched for positive (D) or negative (D’) coefficients of the WGnm_All model.

The modeling process identified genes with positive or negative coefficients with age (Figure 6B). For the model built with the whole-genome expression profile of all samples (WGnm_All), the most significant GO (Gene Ontology) biological process enriched for genes with positive coefficients is GO:0019732 “antifungal humoral response” (Figure 6D). For genes with negative coefficients with age, the enriched biological processes are those involved in mitochondrial energy metabolism (Figure 6D’). This indicates that, as observed in mammalian aging, innate immune genes in fruit flies are positively correlated with aging, while mitochondrial transporters are negatively correlated. Overall, variables with positive coefficients contributed more to the deviance in age than those with negative coefficients (Figure 6B).

To reduce the dimensions of the model, we tried to use the expression values of only immune-defense genes for modeling. The mean MAE was 0.644 (3.22 days) for the model built with all samples. Again, models using only male samples achieved a better mean MAE of 0.424 (2.12 days). Since this model only considers genes that could be directly involved in inflammation, we refer to it as the *Drosophila inflammatory age* (DiAge) model and to values derived from this model as a specimen’s *inflammatory age* or DiAge.

We applied the model to predict the DiAge of both control samples and the high-fat-diet (HFD) samples generated by Stobdan et al. [24]. Interestingly, although flies fed with a normal diet and HFD had the same chronological age, the DiAge of HFD females was significantly higher than those on a normal diet (Figure 7B). This increase in inflammatory age following a high-fat diet was only seen in females but not males. This was still the case even when the profiles were re-evaluated with the male-specific model.

**Figure 7.**
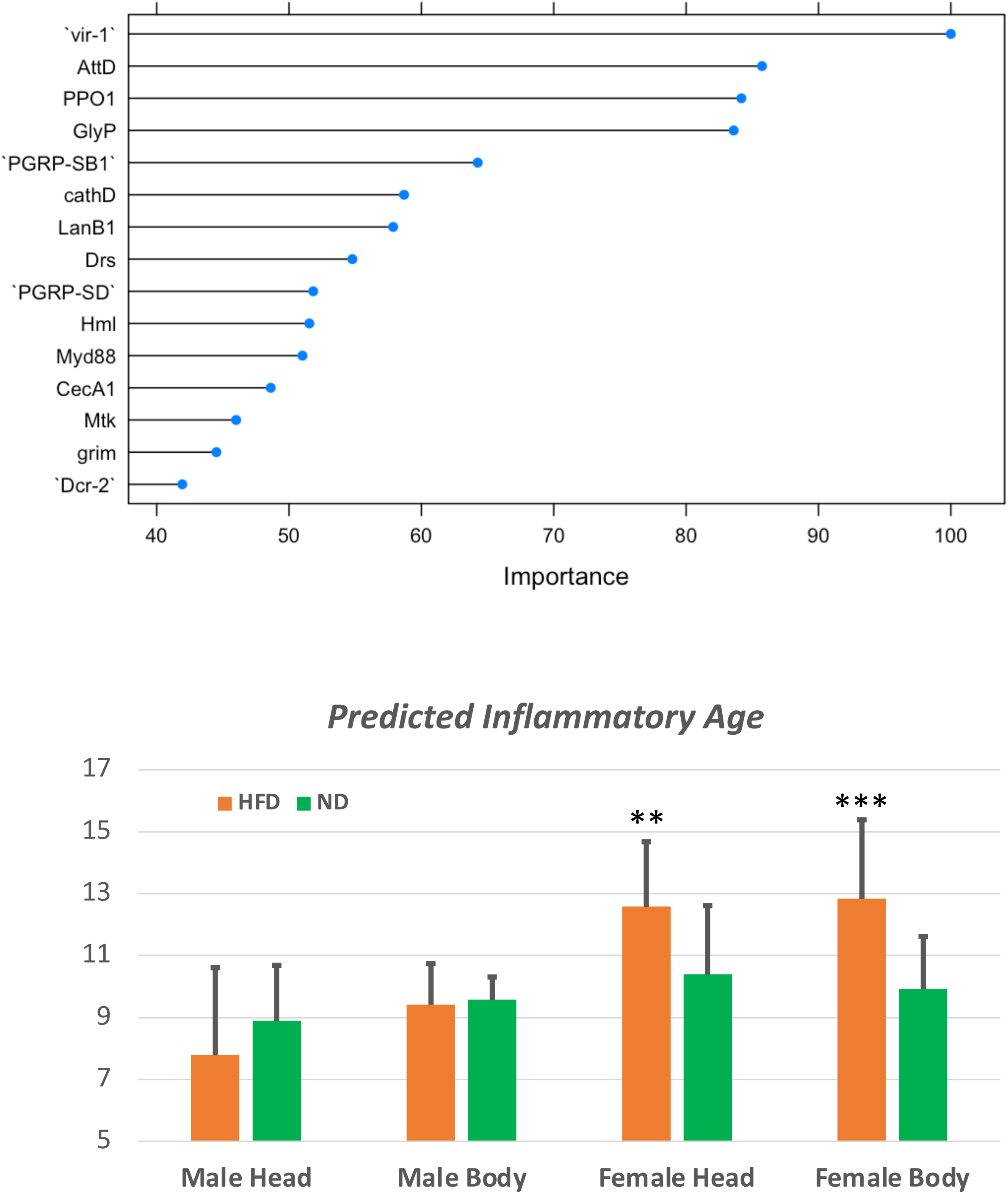
Predicting inflammatory age with the DiAge model. A. The top 15 coefficients of the DiAge model. B. Predicted inflammatory age for male and female flies fed with normal diet (ND) or high-fat diet (HFD). ** p<0.01; *** p<0.001 based on T-test.

## Discussion

Our study demonstrated that a core group of immune-defense genes are consistently upregulated in old age among all strains that were examined so far. These should be considered as markers for future studies of inflammaging in the fruit fly model. Besides the AMP genes, we showed that *SPH93*, a potential immune modulator, also demonstrates prominent age-dependent upregulation. We further showed that it is possible to derive the inflammatory age of the fruit fly using RNA-Seq profiles.

### Reliable inflammaging markers for whole-body samples

The identification of inflammaging markers shared by different fly strains (Figure 1, Table 2) is a surprising finding. Previous studies by different labs have documented aging-associated increases in various AMP genes. However, because the expression of anti-microbial genes is potentially sensitive to pathogens in the environment, it was not clear whether reliable inflammaging patterns could be observed in different settings. Our study indicates that inflammaging at the whole-body level can be reliably detected using the AGE-Index. By focusing on the ratio of the expression levels between old and young flies, instead of the actual expression level, the AGE-Index potentially minimizes the impact of different lab environments and variations of environmental pathogen levels.

On the other hand, the interpretation of AGE-Index values needs to be supplement by actual expression values, as indicated by our analysis of the underlying reasons for the female-skewed patterns.

In light of the possible bias introduced in the mapping and quantification of RNA-Seq data, the AGE-Index values for the public datasets were calculated using the TPM or RPKM values provided by the original studies. Our own RNA-Seq data was processed using a more recent software package. Thus, the shared inflammaging pattern is detectable using different sequencing platforms and appears to hold regardless of the analysis software or pipeline used. We encourage other groups to test this approach further and consider testing the genes we identified as inflammaging markers.

### *SPH93* as a potential inflammaging regulator

We identified novel inflammaging markers such as *SPH93* (*Serine protease homolog 93; CG6639*). The top human ortholog of *SPH93* is *TPSAB1*(*tryptase alpha/beta 1*), a gene implicated in the pathogenesis of asthma and other inflammatory disorders [25,26]. The expression of *SPH93* is responsive to bacterial, viral, or fungal infection (refs). The magnitude of the age-associated increase of *SPH93* is higher than that of most of the AMP genes (Figure 1). Serine proteases, such as *SPE (spz processing enzyme*), are activators of the Toll pathway. Yet a large-scale screening for genes involved in the Toll pathway found that knocking down *SPH93* had no impact on the Toll-mediated response to bacterial infection [27]. Correspondingly, sequence analysis suggests that *SPH93* does not have a functional protease motif [28]. Thus, the functional mechanism of *SPH93* remains to be explored.

The aging dataset of the TM lines highlights the relative importance of genotype or epigenotype in determining inflammaging patterns (Figure 2). There is a clear difference in inflammaging patterns between the TM-O and TM-B lines, presumably the consequence of selecting for postponed senescence. It remains to be tested whether this difference simply reflects the changes in the aging process or is responsible for the extended lifespan of the TM-O lines. A recent analysis by Hanson and Lemaitre [29] has shown that deleting most of the AMP genes did not have a direct impact on aging. Our analysis showed that potential immune modulators, such as *SPH93*, also demonstrate a strong age-associated increase in expression. It remains to be tested whether the inflammaging pattern of *SPH93* is just another phenomenon associated with aging or actively participates in determining the healthspan and lifespan of the animal. It is worth noting that the human serine protease inhibitor AAT suppresses inflammaging and extends both lifespan and healthspan when expressed from the fat body in transgenic fruit flies [10,30]. And this anti-inflammaging function of AAT is independent of its ability to inhibit proteases.

### What tissue to monitor for inflammaging

It is critical to pinpoint which organs are responsible for the inflammaging pattern observed for whole-body samples. However, we found that for tissues dissected from live animals, it is rather challenging to obtain a consistent RNA-Seq or QPCR readout. A previous study reported that inflammaging patterns were observed in samples from the whole gut but not the midgut, which accounts for ∼70–80% of the whole gut [31]. We noticed that the expression profiles of gut samples were strongly influenced by whether and to what extent other organs attached to the gut, such as the Malpighian tubules, were included in the dissected samples. In addition, there was a significant difference in RNA-Seq profiles depending on the operator, even though the same protocol was used. These details all highlight the challenge of monitoring tissue-specific inflammaging patterns via RNA-Seq or QPCR. Gene-specific reporters might be a more suitable alternative for this purpose. In contrast to the guts or fat bodies, heads can be separated from the bodies using metal mesh following snap freezing in liquid nitrogen.

This produces minimal, if any, alteration in the transcriptome during the separation process and seems to give a much more reliable readout of inflammaging patterns.

For whole-body samples, RNA-Seq revealed that the expression levels of inflammaging markers differ significantly between males and females, which may be due to differential expression in sexual organs specifically. Other factors, such as differences in body size, may also confound direct comparison of the sexes. Nonetheless, we found that in both heads and fat bodies, the difference between the sexes increases in old age. This finding is still preliminary given the limited data, yet the observed differences at the whole-body and organ-specific levels warrant the two sexes being monitored separately in any aging analysis.

### Predicting Inflammatory Age with the DiAge model

In this work, we tried to extrapolate to the fruit fly the kind of elastic-net model used by Horvath to determine the biological age of human samples [11]. For this purpose, gene expression value was min-max normalized to mimic the range of DNA methylation data. We showed that it is feasible to measure the inflammatory age of *Drosophila* (DiAge) by assessing head or body RNA-Seq profiles with the model built with normal expression profiles. The current DiAge model was based on a combination of the *w1118*, *Canton-S-w1118*, and *modENCODE* (*y^1^; cn bw^1^sp^1^*) strains due to the need to accumulate enough data. Based on the standard resampling cross-validation, the median absolute error (MAE) of the model was ∼3.35 days, or roughly equivalent to 3.89 human years. This could be significantly improved by training the model with additional datasets, especially expression profiles for more advanced ages. The accuracy of the model should be further improved by building individual ones for specific strains, such as the *w1118* or *Canton-S,* as we noticed that including or excluding data from the *wDah* strain had a significant impact on the mean MAE of the model. With the increasingly lower cost of high-throughput sequencing, we anticipate more adaptation of RNA-Seq as a standard measurement of inflammaging and are looking forward to improving the model in collaboration with the community.

For aging studies in *Drosophila*, monitoring lifespan is and should be the gold standard. However, a model based on RNA-Seq can be used to study the dynamics of biological aging following certain environmental or genetic events. Our model predicted that the high-fat diet (20% coconut oil) significantly accelerated inflammaging in females but not males based on the expression profiles generated by Stobdan et al. [24] (Figure 8B). It needs to be emphasized that this particular diet experiment was conducted on the *w1118* strain, which accounts for the majority of the normal expression profiles used to build the predictive model. Interestingly, a separate study found that a similar high-fat diet regime (10% or 30% coconut oil) shortened the lifespan of *w1118* females [32]. A surprise from the model prediction was that, unlike the female, male flies did not demonstrate the detectable acceleration of inflammaging. A more recent study, published after we had done the initial machine learning and prediction, showed that the impact of a high-fat diet on body weight and composition is specific to female *w1118* flies but not to males [33]. In fact, another recent study reported that a high-fat diet actually increased the lifespan of *w1118* males [34]. Even though the fruit fly has a much shorter lifespan compared with mammalian models, monitoring survival over 70 days is still cumbersome and subject to disruption by many uncontrolled variables. Our DiAge model promises an alternative measurement that could be applied at any point following a particular treatment to measure the impact on inflammaging.

## Detailed Methods

### Calculating AGE-Index values

To take advantage of the vast amount of RNA-Seq data available in the public domain, we developed a simple method to identify genes whose expression levels are significantly increased in older fruit flies. For data sets that have samples of both young and old adults, we assign the <u>a</u>ging-associated <u>g</u>ene <u>e</u>xpression index (AGE-Index) for each gene as the ratio of its expression level at the old stage divided by that at the young stage. We adopt a norm that has been widely used in the field: adults at 3 to 7 days following eclosion are considered as in the reference young age. Reciprocally, adults aged 30 days or more after eclosion are considered as entered the “old” stage. This 30-day threshold is somewhat arbitrary and may not apply equally to different strains, which we will elaborate on in the following sections.

Unless otherwise mentioned, we use the TPM (transcripts <u>p</u>er <u>m</u>illion) value derived from the RNA-Seq data to represent the expression levels of every gene in a given sample. We compared the relationship between the AGE-Index calculated based on RNA-Seq data and the folder changes detected using SYBR-green-based quantitative PCR. There is a linear correlation between the two, even though the ratio of the two is not 1:1 (Figure 1). We also examined the potential difference of using RPKM or FPKM instead of TPM. We found that for the samples we tested, the difference was minor.

Since the total sum of expression values for each sample adds to 1,000,000 with TPM, it may be more suitable for cross-sample comparison.

During our preliminary analysis, we found the expression levels of some innate immune genes differed significantly between male and female adults of the same age. Thus, AGE-Index is always calculated separately for males and females. RNA-Seq datasets without clear information on the sex of the samples were excluded from this analysis.

### Datasets for identifying shared inflammaging patterns

We used four datasets for this purpose. The first was the *modENCODE* developmental stage dataset [19], based on multiple samples collected and processed at different times, with some generated from different labs. The isogenic *y^1^; cn bw^1^sp^1^* strain used for this study served as the background strain for the *D. melanogaster* genome project [35]. The normalized gene expression value in this dataset is used by FlyBase (www.flybase.org) to represent the developmental expression pattern of each gene. We calculated the AGE-Index for this dataset using the expression level of day 30 divided by that of day 5.

The second dataset was generated by Trudy Mackay’s group [16]. This set of lines originated from an outbred of flies collected around Amherst, MA [36]. Five baselines (*TM-B*) had been reared separately for over 30 years (850 generations) and served as controls for a postponed senescence evolution experiment. In addition, five O lines (*TM-O*) were selected for postponed senescence and had been reared separately for over 170 generations at the time of the RNA-Seq experiment. On average, the lifespan of the TM-O lines is about 70% longer than that of the *TM-B* lines [16,36]. For the total of 10 lines (5 B lines and 5 O lines), gene expression profiles were obtained at week 1 (days 3–5) or week 5 (days 33–35) for both males and females. For each line/sex, two replicated RNA samples were processed for RNA-Seq. We calculated the AGE-Index value for the individual lines using the expression profile data included as a supplement table [16] (SF3)

Two RNA-Seq datasets were generated by us using *Oregon-R* and *w1118* strains, respectively. *Oregon-R* is one of the most often used wild-type strains, and *w1118* is widely used as the starting strain for transposon insertions and other mutagenesis projects. Male and female flies were separated 1 day after eclosion and aged to day 5, 30, 45 in 25°C. 2-3 samples for each sex/age combination were processed for RNA-Seq.

### Datasets for organ-specific inflammaging

The *GSE130158* dataset has expression profiles for the brain, thorax, fat body, and gut on days 10, 30, and 50. Only samples from females were included in the original study. The *white Dahomey* (*wDah*) strain is known to carry the endosymbiotic bacterium *Wolbachia*, which interacts with the insulin/IGF signaling pathway [37].

*GSE110349* was originally used to monitor the changes in intron retention during aging [23]. It contains RNA-Seq data for the heads of *w1118* males on days 10, 20, 30, and 50. We used the expression value on day ten as the reference to calculate the AGE-Index for each gene.

Using the *w1118* strain (RRID: BDSC_5905), we generated RNA-Seq profiles for dissected male and female fat body and gut on days 5, 30, and 45. Expression levels on day five were used as a reference to calculate AGE-Index values for day 30 and day 45.

### Datasets for machine learning

Our analysis of data generated by different labs indicated that RNA samples obtained from the whole body or heads had the most reliable and repeatable inflammaging patterns. In contrast, RNA samples involving the dissection of live animals showed significant batch differences in terms of innate immune gene expression. In light of this observation, we compiled a dataset for machine learning by searching for RNA-Seq profiles derived from the whole body, head, or body without head (S7).

Samples with clear age and growth condition (25 °C) information were collected. Wherever possible, age information was verified by checking the publication associated with the dataset. The raw read files were then downloaded from SRA at NCBI. Transcripts per million (TPM) value for every gene was obtained using Salmon [17].

Wild-type flies raised at 25 °C with standard food were considered “normal” samples and was used for generating progression models using an elastic net.

The majority of the RNA-Seq samples we identified were used as controls in 17 different studies, most of which did not focus on aging. Wild-type strains such as *w1118* and *Canton-S* are often used as controls; therefore, samples from these two accounted for most of the learning set. This selection process resulted in 487 RNA-Seq profiles, of which 175 were from female samples and 236 were from male samples. Another 26 profiles were derived from aged adults of mixed sexes. The remaining 50 profiles do not have identifiable information on the sex identity of the samples.

For all of these samples, we downloaded and processed the raw reads and obtained TPM for each sample using Salmon. We subjected this dataset to elastic-net-based machine learning using the glmnet r package [18]. Unless otherwise specified, the sample set was randomly split 0.7:0.3 for training and testing, respectively. We tried different normalization methods and chose to proceed with the min-max normalization. For each gene, its expression level in sample x is transformed to (TPMx – TPMmin) / (TPMmax – TPMmin). Inspired by the work of Horvath [11], we transformed the age of the samples with T_Age=(age – 1)/5.

### Dataset for high-fat diet

Stobdan et al. conducted an experiment whereby the 2–3 day old male and female flies were fed seven days with a normal diet (ND) or high-fat diet (HFD 20% coconut oil) [24]. Immediately following the treatment period, the fruit flies were processed for RNA-Seq. The study found that male and female flies had distinct transcriptomic responses to a high-fat diet. However, the impact of a high-fat diet on aging was not assessed in the original study. The control samples produced by this study, those fed with a normal diet, were included in the learning dataset for building the model. Based on how the samples were aged, we estimated the average age of samples to be 9.5 days. Expression profiles from samples fed with a high-fat diet were not used to build the model.

### Immune-defense genes

To focus our attention on the genes involved in immunity and defense against pathogens, we compiled a list based on the annotation recorded by the FlyBase [38]. Using AmiGO2, we identified 322 genes associated with GO:0002376, “immune system process,” and 364 genes associated with GO:0098542, “defense response to other organisms”. The Immune-defense (ImmDef) gene list is a fusion of the above two lists and contains 504 genes, accounting for about 3.7% of the genome. Unless otherwise mentioned, ImmDef genes are marked as solid red dots in AGE-Index plots (Figure 2).

### Fly stocks used in this study

We generated our own aging RNA-Seq datasets with the *Oregon-R* (RRID: BDSC_6363) strain and the *w1118* (RRID: BDSC_5905) strain. The BDSC_5905 line has been isogenized and tested for normal learning, memory, and circadian rhythms [39]. Our copies of both strains were obtained from the Bloomington stock center around 2010 and maintained in the lab under standard conditions.

### RNA Extraction and qPCR

The culture and aging of *Drosophila* are carried out as previously described [10]. Briefly, fruit flies were maintained and aged in a 25 °C incubator with a 12:12 light: dark cycle. Eclosion is counted as day 1. RNA was extracted with Trizol, and cDNA was synthesized immediately following RNA extraction using the High-Capacity cDNA Reverse Transcription Kit (Fisher Cat # 4368814) that was originally developed by Applied Biosystems. QPCR was carried out with SYBR Green PCR Mix (Applied Biosystems Cat # 4309155) using an ABI 7500 Real-Time PCR System.

### RNA-Seq and data analysis

RNA samples with RIN greater than eight were processed for library construction by the UF ICBR following standard Illumina protocol. Sequencing was done with Illumina HiSeq 2500 or NovaSeq 6000. Following the removal of low-quality reads and adaptor sequences, the reads were mapped to the Dm6.43 genome with STAR [40] and used for CUFFDIFF analysis [20]. A parallel pipeline consisting of Hisat2 and stringtie [41] was applied to identify differentially expressed genes with DESeq2 [42].

## References

1. López-Otín C, Blasco MA, Partridge L, Serrano M, Kroemer G. The hallmarks of aging. Cell. 2013;153:1194–217.

2. de Magalhães JP, Curado J, Church GM. Meta-analysis of age-related gene expression profiles identifies common signatures of aging. Bioinformatics. 2009;25:875– 81.

3. Frenk S, Houseley J. Gene expression hallmarks of cellular ageing. Biogerontology. Springer Netherlands; 2018;19:547–66.

4. Fushan AA, Turanov AA, Lee SG, Kim EB, Lobanov A V., Yim SH, et al. Gene expression defines natural changes in mammalian lifespan. Aging Cell. 2015;14:352– 65.

5. Tyshkovskiy A, Ma S, Shindyapina A V., Tikhonov S, Lee SG, Bozaykut P, et al. Distinct longevity mechanisms across and within species and their association with aging. Cell. 2023;186:2929–2949.e20.

6. Ferrucci L, Fabbri E. Inflammageing: chronic inflammation in ageing, cardiovascular disease, and frailty. Nat. Rev. Cardiol. Springer US; 2018;15:505–22.

7. Lemaitre B, Hoffmann J. The host defense of Drosophila melanogaster. Annu. Rev. Immunol. 2007;25:697–743.

8. Buchon N, Silverman N, Cherry S. Immunity in Drosophila melanogaster — from microbial recognition to whole-organism physiology. Nat. Rev. Immunol. Nature Publishing Group; 2014;14:796–810.

9. Meekins DA, Kanost MR, Michel K. Serpins in arthropod biology. Semin. Cell Dev. Biol. Elsevier Ltd; 2017;62:105–19.

10. Yuan Y, DiCiaccio B, Li Y, Elshikha AS, Titov D, Brenner B, et al. Anti-inflammaging effects of human alpha-1 antitrypsin. Aging Cell. 2018;17:1–11.

11. Horvath S. DNA methylation age of human tissues and cell types. Genome Biol. 2015;16.

12. Sayed N, Huang Y, Nguyen K, Krejciova-Rajaniemi Z, Grawe AP, Gao T, et al. An inflammatory aging clock (iAge) based on deep learning tracks multimorbidity, immunosenescence, frailty and cardiovascular aging. Nat. Aging. 2021;1:748–748.

13. Peters MJ, Joehanes R, Pilling LC, Schurmann C, Conneely KN, Powell J, et al. The transcriptional landscape of age in human peripheral blood. Nat. Commun. 2015;6.

14. Pletcher SD, Macdonald SJ, Marguerie R, Certa U, Stearns SC, Goldstein DB, et al. Genome-wide transcript profiles in aging and calorically restricted Drosophila melanogaster. Curr. Biol. 2002;12:712–23.

15. Landis GN, Abdueva D, Skvortsov D, Yang J, Rabin BE, Carrick J, et al. Similar gene expression patterns characterize aging and oxidative stress in Drosophila melanogaster. Proc. Natl. Acad. Sci. U. S. A. 2004;101:7663–8.

16. Carnes MU, Campbell T, Huang W, Butler DG, Carbone MA, Duncan LH, et al. The genomic basis of postponed senescence in Drosophila melanogaster. PLoS One. 2015;10:1–22.

17. Patro R, Duggal G, Love MI, Irizarry RA, Kingsford C. Salmon provides fast and bias-aware quantification of transcript expression. Nat. Methods. Nature Publishing Group; 2017;14:417–9.

18. Friedman J, Hastie T, Rob Tibshirani. Regularization Paths for Generalized Linear Models via Coordinate Descent. J. Stat. Softw. 2010;33:1–22.

19. Consortium T, Roy S, Ernst J, Kharchenko P V, Kheradpour P, Negre N, et al. Identification of Functional Elements and Regulatory Circuits by Drosophila modENCODE. Science (80-.). 2011;330:1787–97.

20. Trapnell C, Roberts A, Goff L, Pertea G, Kim D, Kelley DR, et al. Differential gene and transcript expression analysis of RNA-seq experiments with TopHat and Cufflinks. Nat. Protoc. Nature Publishing Group; 2012;7:562–78.

21. Junell A, Uvell H, Davis MM, Edlundh-Rose E, Antonsson Å, Pick L, et al. The POU Transcription Factor Drifter/Ventral veinless Regulates Expression of Drosophila Immune Defense Genes . Mol. Cell. Biol. 2010;30:3672–84.

22. Flybase. IM18 [Internet]. [cited 2024 Jan 3]. Available from: https://flybase.org/reports/FBgn0067903#expression

23. Adusumalli S, Ngian ZK, Lin WQ, Benoukraf T, Ong CT. Increased intron retention is a post-transcriptional signature associated with progressive aging and Alzheimer’s disease. Aging Cell. 2019;18:1–13.

24. Stobdan T, Sahoo D, Azad P, Hartley I, Heinrichsen E, Zhou D, et al. High fat diet induces sex-specific differential gene expression in Drosophila melanogaster. PLoS One. 2019;14:1–19.

25. Lyons JJ, Chovanec J, Connell MPO, Liu Y, Selb J, Schwartz LB, et al. Heritable risk for severe anaphylaxis associated with increased a -tryptase – encoding germline copy number at TPSAB1. 2020;622–32.

26. Greiner G, Sprinzl B, Górska A, Ratzinger F, Gurbisz M, Witzeneder N, et al. Hereditary α tryptasemia is a valid genetic biomarker for severe mediator-related symptoms in mastocytosis. Blood. 2021;137:238–47.

27. Kambris Z, Brun S, Jang IH, Nam HJ, Romeo Y, Takahashi K, et al. Drosophila Immunity: A Large-Scale In Vivo RNAi Screen Identifies Five Serine Proteases Required for Toll Activation. Curr. Biol. 2006;16:808–13.

28. Cao X, Jiang H. Building a platform for predicting functions of serine protease-related proteins in Drosophila melanogaster and other insects. Insect Biochem. Mol. Biol. Elsevier; 2018;103:53–69.

29. Hanson MA, Lemaitre B. Antimicrobial peptides do not directly contribute to aging in Drosophila, but improve lifespan by preventing dysbiosis. DMM Dis. Model. Mech. 2023;16.

30. Yuan Y, Van Belkum M, O’Brien A, Garcia A, Troncoso K, Elshikha AS, et al. Human Alpha 1 Antitrypsin Suppresses NF-κB Activity and Extends Lifespan in Adult Drosophila. Biomolecules. 2022;12:1–16.

31. Chen H, Zheng X, Zheng Y. Age-associated loss of lamin-b leads to systemic inflammation and gut hyperplasia. Cell. Elsevier; 2014;159:829–43.

32. Liao S, Amcoff M, Nässel DR. Impact of high-fat diet on lifespan, metabolism, fecundity and behavioral senescence in Drosophila. Insect Biochem. Mol. Biol. 2021;133.

33. Eickelberg V, Rimbach G, Seidler Y, Hasler M, Staats S, Lüersen K. Fat Quality Impacts the Effect of a High-Fat Diet on the Fatty Acid Profile, Life History Traits and Gene Expression in Drosophila melanogaster. Cells. 2022;11.

34. Shi D, Han TS, Chu X, Lu H, Yang X, Zi TQ, et al. An isocaloric moderately high-fat diet extends lifespan in male rats and Drosophila. Cell Metab. Elsevier Inc.; 2021;33:581–597.e9.

35. Celniker SE, Wheeler DA, Kronmiller B, Carlson JW, Halpern A, Patel S, et al. Finishing a whole-genome shotgun: release 3 of the Drosophila melanogaster euchromatic genome sequence. Genome Biol. 2002;3:1–14.

36. Rose MR. Laboratory Evolution of Postponed Senescence in Drosophila melanogaster. Evolution (N. Y). 1984;38:1004.

37. Grönke S, Clarke D-F, Broughton S, Andrews TD, Partridge L. Molecular evolution and functional characterization of Drosophila insulin-like peptides. PLoS Genet. 2010;6:e1000857.

38. Gramates LS, Agapite J, Attrill H, Calvi BR, Crosby MA, dos Santos G, et al. FlyBase: a guided tour of highlighted features. Genetics. 2022;220.

39. Ryder E, Blows F, Ashburner M, Bautista-Llacer R, Coulson D, Drummond J, et al. The DrosDel collection: A set of P-element insertions for generating custom chromosomal aberrations in Drosophila melanogaster. Genetics. 2004;167:797–813.

40. Dobin A, Davis CA, Schlesinger F, Drenkow J, Zaleski C, Jha S, et al. STAR: Ultrafast universal RNA-seq aligner. Bioinformatics. 2013;29:15–21.

41. Kim D, Pertea M, Kim D, Pertea GM, Leek JT, Salzberg SL. Transcript-level expression analysis of RNA-seq experiments with HISAT, StringTie and Ballgown. Nat. Protoc. Nature Publishing Group; 2016;11:1650–67.

42. Anders S, McCarthy DJ, Chen Y, Okoniewski M, Smyth GK, Huber W, et al. Count-based differential expression analysis of RNA sequencing data using R and Bioconductor. Nat. Protoc. 2013;8:1765–86.

