## supplements for "Shared Transcriptomic Signatures of Inflammaging Among Diverse Strains of *Drosophila melanogaster*"

List of Supplementary files:

SF1. Correlation between AGE_Index values and folder change detected by QPCR.

SF2. Full range plots for Figure 1

SF3. log2(AGE-Index) plots for all immune-defense genes.

SF4. AGE_Index plot for individual TM lines.

*S1. Table of AGE_Index values for the whole-body datasets.*
*AGE_Index* values were calculated for the 5 whole-body datasets.

*S2. Significantly changed genes in day 30 Oregon-R.*
Significantly changed genes identified by DESeq2 for day-30 male vs. day-5 male, and day-30 female vs. day-5 female.

*S3. Significantly changed genes in day 30 and day 45 of w1118.*
Significantly changed genes identified by DESeq2 or CuffDiff for day-30 and day-45 male vs. day-5 male, and day-30 and day-45 female vs. day-5 female.

*S4. Table of AGE_Index for organs and body parts of wDah females (GSE130158).*

*S5. Table of AGE_Index for heads of w1118 male (GSE110349).*

*S6. Table of AGE_Index for fatbody and guts of w1118 male and female (this study).*

*S7. Machine-learning dataset used for DiAge training and testing.*

*S8. Genes and their coefficients in the corresponding models.*

*S9. GO enrichment analysis of genes with positive or negative coefficients.*
